# Mapping the substrate sequence and length of the *Plasmodium* M1 and M17 aminopeptidases

**DOI:** 10.1101/2020.10.13.338178

**Authors:** Tess R Malcolm, Karolina W. Swiderska, Brooke K Hayes, Marcin Drag, Nyssa Drinkwater, Sheena McGowan

**Affiliations:** Monash Biomedicine Discovery Institute and Department of Microbiology, Monash University, Clayton, 3800, Australia; Department of Chemical Biology and Bioimaging, Wroclaw University of Science and Technology, Wyb. Wyspianskiego 27, 50-370 Wroclaw, Poland

**Author notes:** Corresponding author: Sheena McGowan.

**Keywords:** leucine aminopeptidase, alanine aminopeptidase, *Plasmodium*, substrate specificity, cooperativity

## Abstract

During malarial infection, *Plasmodium* parasites digest human hemoglobin to obtain free amino acids for protein production and maintenance of osmotic pressure. The *Plasmodium* M1 and M17 aminopeptidases are both postulated to have an essential role in the terminal stages of the hemoglobin digestion process and are validated drug targets for the design of new dual-target anti-malarial compounds. In this study, we profiled the substrate specificity fingerprints and kinetic behaviors of M1 and M17 aminopeptidases from *Plasmodium falciparum* and *Plasmodium vivax,* and the mouse model species, *Plasmodium berghei.* We found that although the *Plasmodium* M1 aminopeptidases share a largely similar, broad specificity at the P1 position, the *P. falciparum* M1 displays the greatest diversity in specificity and *P. berghei* M1 showing a preference for charged P1 residues. In contrast, the *Plasmodium* M17 aminopeptidases share a highly conserved preference for hydrophobic residues at the P1 position. The aminopeptidases also demonstrated intra-peptide sequence specificity, particularly the M1 aminopeptidases, which showed a definitive preference for peptides with fewer negatively charged intrapeptide residues. When tested with a panel of peptides of increasing length, each aminopeptidase exhibited unique catalytic behavioral responses to the increase in peptide length, although all six aminopeptidases exhibited an increase in cooperativity as peptide length increased. Overall the *P. vivax* and *P. berghei* enzymes were generally faster than the *P. falciparum* enzymes, which we postulate is due to subtle differences in structural dynamicity. Together, these results build a kinetic profile that allows us to better understand the catalytic nuances of the M1 and M17 aminopeptidases from different *Plasmodium* species.

## Introduction

Malaria is one of the world’s most prevalent parasitic diseases, mostly commonly caused by *Plasmodium falciparum* and *Plasmodium vivax.* During the blood stage of infection, *Plasmodium* parasites employ a cascade of proteases to digest hemoglobin to liberate amino acids for use in protein production and maintenance of osmotic pressure within the red blood cell (1). The alanyl M1 and leucine M17 metalloaminopeptidases from *P. falciparum, Pf*A-M1 and *Pf*A-M17, are postulated to play an essential role in the terminal stages of this hemoglobin digestion pathway (2–5). Attempts to knockout the genes as well as inhibition studies both *in vitro* and *in vivo* have suggested that these proteases are essential and non-redundant (2, 4, 6, 7). In our laboratory, *Pf*A-M1 and *Pf*A-M17 are the subject of an ongoing structure-based drug design study aimed at generating new non-artemisinin based anti-malarial agents (8–11). This program has been successful in developing small molecule dual inhibitors of *Pf*A-M1 and *Pf*A-M17 that also show anti-parasitic activity. As anti-malarial resistance becomes more widespread, a requirement for future antimalarials is cross-species activity to ensure that new drugs are effective against both falciparum and vivax malaria (12). Our latest generation of compounds meet this criterium and possess potent inhibition of the *Plasmodium vivax* M1 and M17 aminopeptidases (*Pv*-M1 and *Pv*-M17). Therefore, the continued development of this dual-target series is complex and demands detailed knowledge about the structure and specificity of each of these targets.

The extended substrate profiles of the *Plasmodium* alanyl M1 and leucyl M17 aminopeptidases have not been investigated to date. The N-terminal, or P1, substrate specificity of both *Pf*A-M1 and *Pf*A-M17 has been profiled, and both possess a broader substrate specificity range at the P1 position than their names suggest (13). *Pf*A-M1 has a broad substrate specificity, processing hydrophobic, polar and non-polar residues alike, whereas *Pf*A-M17 has a comparatively narrow substrate specificity profile, with a distinct preference for large hydrophobic residues at the P1 position. Further analysis of the *Pf*A-M1 substrate specificity profile has been carried out, extending into the P1’ and P2’ pockets (14). *Pf*A-M1 has the highest affinity for di-peptide substrates with large residues at the P1’ position and a comparatively low affinity for di-peptide substrates with small residues at the P1’ position (14). The P2’ position also favors large, hydrophobic residues (14) and when processing tri-peptides, N-terminal residues are liberated singularly in a processive fashion (15). Beyond the hydrolysis of tripeptides, there is evidence that suggest both M1 and M17 aminopeptidases are active against long peptide substrates. *Pf*A-M1 has been shown to digest the hemoglobin-derived hexapeptide VDPENF (4) which is not unusual for the enzyme family and the human M1 aminopeptidase ERAP1 has been shown to hydrolyze peptide substrates up to 12 residues long (16). While *Pf*A-M17 has not been directly shown to hydrolyze long peptide substrates, the M17 aminopeptidase from tomatoes (LAP-A) has been suggested to have activity against a 19-residue precursor to the peptide systemin (17, 18).

Here, we describe the substrate profiles for the M1 and M17 aminopeptidases from the malaria causing *P. falciparum* (*Pf*A-M1 and *Pf*A-M17), *P. vivax* (*Pv*-M1 and *Pv*-M17), as well as the parasite used in the murine malaria models, *Plasmodium berghei* (*Pb*-M1 and *Pb*-M17). This study thereby characterizes the substrate profiles of the key *Plasmodium* clinical and animal model species and will inform pre-clinical development of novel inhibitor compound series. The results demonstrate that the activity profiles of the M1 and M17 *Plasmodium* homologs are not only influenced by the N-terminal/P1 residue, but also by substrate length and intrapeptide sequence. This study will underpin our understanding of how substrate characteristics dictate *Plasmodium* M1 and M17 catalytic behavior.

## Materials and Methods

### Reagents

Genes encoding *Pb*-M1 (PlasmoDB ID: PBANKA_1410300, UniProt ID: A0A509ATR4) and *Pb*-M17 (PlasmoDB ID: PBANKA_1309900, UniProt ID A0A509APA2) were purchased from GenScript. The synthesized genes were codon optimized for gene expression in *Escherichia coli,* encoded an in-frame C-terminal hexa-histidine tag and were cloned into pET-21d expression vectors. All peptide substrates were purchased from GL Biochem and purified by HPLC to >95% by MS analysis and chemicals purchased from commercial suppliers.

### Production and Purification of Recombinant Aminopeptidases

Recombinant *Pf*A-M1 (UniProt ID O96935), *Pf*A-M17 (UniProt ID Q8IL11), *Pv*-M1 (UniProt ID A5JZA9) and *Pv*-M17 (UniProt ID A5K3U9) proteins were produced from constructs described previously (11). Recombinant *Pb*-M1 and *Pb*-M17 were purified using a similar protocol where recombinant protein was expressed in *E. coli* BL31 DE3 cells. Cells were grown in auto-induction media to force overexpression of target proteins before *E. coli* cells were lysed by sonication, centrifuged and the supernatant pooled. Protein was purified via nickel-affinity chromatography, which utilized the encoded C-terminal hexa-histidine tag (HisTrap™ Ni^2+^-NTA column; GE Healthcare Life Sciences), and size-exclusion chromatography (Superdex S20010/300 column; GE Healthcare Life Sciences). All aminopeptidases were stored at −80°C until use; M1 aminopeptidases were stored in 50 mM HEPES pH 8.0, 0.3 M NaCl and 5% glycerol, and M17 aminopeptidases were stored in 50 mM HEPES pH8.0 and 0.3 M NaCl.

### Enzyme Kinetics Using Fluorogenic Substrates

Aminopeptidase enzyme assays were carried out in white 384-well plates (Axygen) in a final volume of 50 μL. Catalytic activity was determined by continual measurement of the liberation of fluorogenic leaving group, 7-amido-4-methyl-coumarin (NHMec) from the commercially available substrate Leucine-7-amido-4-methyl-coumarin (Sigma-Aldrich). For processivity assays, synthetic peptide substrates Leucine-Leucine-7-amido-4-methyl-coumarin and Leucine-Leucine-Leucine-Leucine-7-amido-4-methyl-coumarin were used. Fluorescence was detected using a FLUOStar Optima plate read (BMG Labtech) with excitation and emission wavelengths of 355 nm and 460 nm respectively. Assays were carried out in 100 mM Tris pH 8.0 and M17 aminopeptidase assays were supplemented with 1 mM CoCl_2_ unless otherwise stated. Assays screening activity in metal ions included CoCl_2_, MnCl_2_, ZnCl_2_ and MgCl_2_ at varied concentrations between 0 and 1000 μM. Final enzyme concentration for each aminopeptidase was *Pf*A-M1 = 20 nM; *Pf*A-M17 = 125 nM; *Pv*-M1 = 10 nM; *Pv*-M17 = 50 nM; *Pb*-M1 = 20 nM; *Pb*-M17 = 100 nM. For calculation of Michaelis-Menten constants enzyme was kept at a constant concentration and substrate titrated from 0 – 500 μM. Assays were carried out in biological triplicate and data was analyzed using PRISM GraphPad 8. Michaelis Menten *K*m, *kcat* and Hill’s coefficient were calculated using the PRISM GraphPad 8 software and presented as mean ± standard error of the mean (SEM).

Substrate-profiling at the P1 position was achieved by the use of a fluorogenic substrate library containing 19 natural amino acids and 49 unnatural amino acids (13). Amino acids were tagged with 7-amino-4-carbamoylmethylcoumarin (ACC) fluorogenic leaving group, which fluoresced upon liberation. Final screening of the library was carried out at 10 μM substrate and 0.50 nM enzyme for M1 aminopeptidase assays, and 500 nM substrate and 6.0 nM enzyme for M17 aminopeptidase assays. Assays were carried out in 50 mM Tris pH 8.0, and M17 aminopeptidase assays were supplemented with 1 mM CoCl_2_. Release of free ACC fluorophore was monitored. For ACC substrates, *K*_m_ values of selected substrates were measured using a range of substrate concentrations unique to each enzyme (*Pv*-M1 = 300 μM – 17.6 μM; *Pb*-M1 = 180 μM – 10.5 μM; *Pv*-M17, *Pb*-M17 = 10 μM – 0.58 μM) and keeping enzyme concentration constant: *Pv*-M1 = 0.672 nM; *Pb*-M1 = 0.336 nM; *Pv*-M17 = *Pb*-M17 = 1.64 nM. Each experiment was repeated at least three times and the average value with standard error was calculated. Concentration of DMSO in the assay was less than 2% (v/v).

### Activity Towards Peptide Substrates

Unlabeled peptide substrates (500 μM) were incubated with enzymes (1 mg/mL) for either 45 mins (*Pf*A-M1, *Pf*A-M17) or 10 mins (*Pv*-M17, *Pb*-M1 & *Pb*-M17) at 37°C in a final reaction volume of 10 μL. *Pv*-M1 cleaved peptides with high efficiency and as such enzyme concentration was decreased to 0.1 mg/mL, peptide concentration reduced to 250 μM and reaction time to 5 mins. Assays were carried out in 50 mM Tris pH 8.0 and M17 aminopeptidase assays were supplemented with 1.0 mM CoCl_2_. The reactions were stopped by heating samples at 100°C for 10 mins. Samples were analyzed via mass spectrometry using ESI (MicroTOFq; Bruker Daltonics). Data was analyzed using Data Analysis V3.4 Flex Analysis V1.4 (Bruker). Relative percentage of substrate cleaved was calculated by integrating the area under the peptide curve. Experiments were carried out in technical duplicate and percentage peptide cut values were graphed using GraphPad PRISM 8.

## Results

### Characterization of recombinant M1 and M17 aminopeptidases from P. berghei

The *Pb*-M1 and -M17 aminopeptidases were successfully expressed using a bacterial expression system. Both *Pb*-M1 and *Pb*-M17 eluted as a single peak at the same elution volumes as their respective *P. falciparum* and *P. vivax* homologs (Fig 1A, 1B), although the presence of a small tail to the right of the main *Pb*-M17 peak indicates there was a low concentration of smaller *Pb*-M17 species (Fig 1B). It is not uncommon for M17 family enzymes to display a mix of oligomeric states in the purified sample (Fig. 1B). The molecular weight and purity were confirmed using SDS-PAGE and showed that each homolog sample was homogenous and of the expected molecular weight (Fig 1A, 1B).

**Figure 1.**
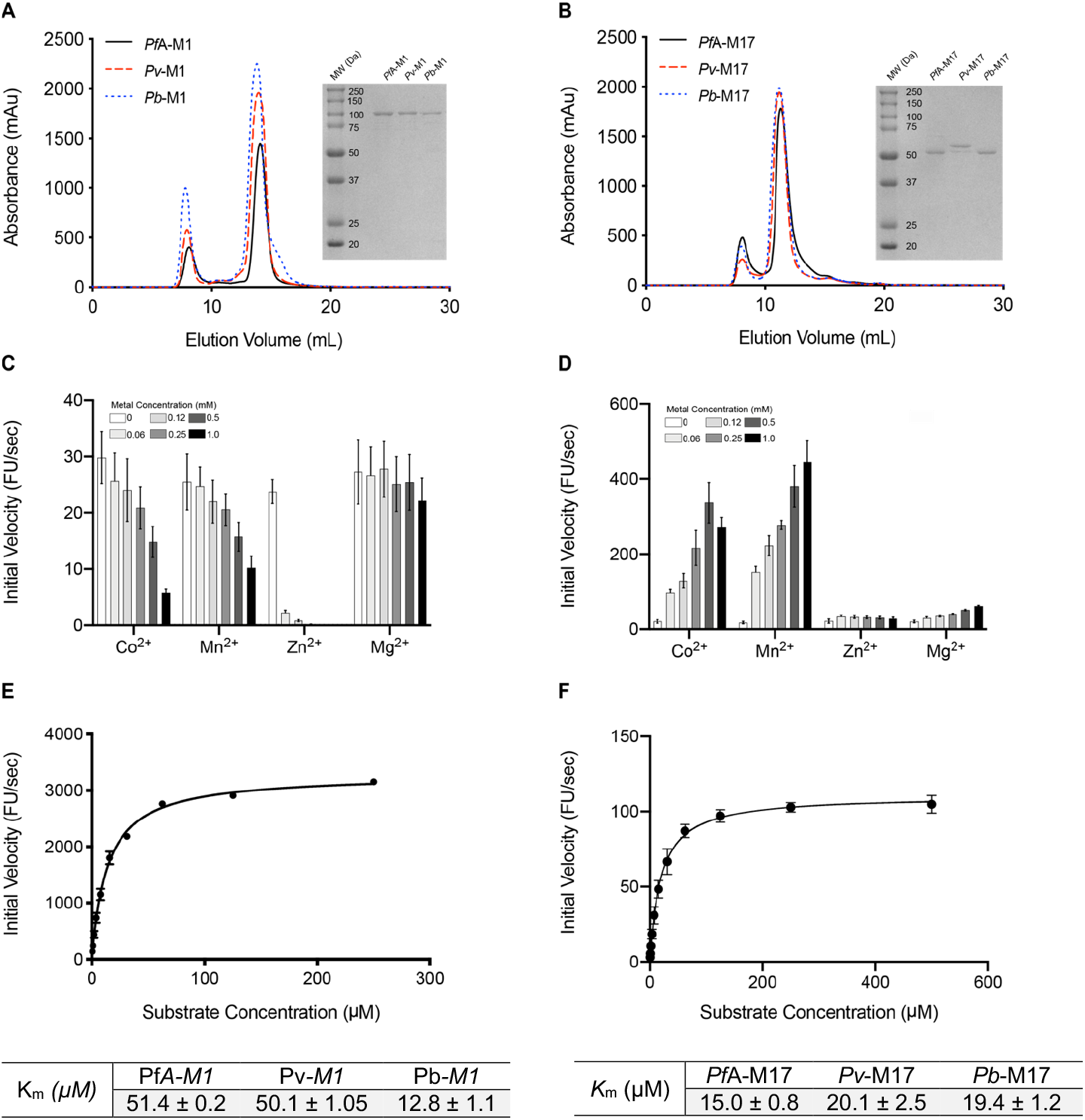
Purification and kinetic characterization of *Plasmodium berghei* M1 and M17 aminopeptidases, *Pb*-M1 and *Pb*-M17. **(A)** Purification profile of *Pf*A-M1 (black), *Pv*-M1 (red dash) and *Pb*-M1 (blue dash) and SDS-PAGE gel (inset) showing correct molecular weight and purity for each homolog. **(B)** Purification profile of *Pf*A-M17 (black), *Pv*-M17 (red dash) and *Pb*-M17 (blue dash) and SDS-PAGE gel (inset) showing correct molecular weight and purity. **(C)** *Pb*-M1 activity profile in presence of increasing concentrations of divalent cations. Activity is highest in the absence of metal ions. **(D)** *Pb*-M17 activity in presence of increasing concentrations of divalent cations. Activity is highest in Co^2+^ and Mn^2+^ and increases as ion concentration increases. **(E)** Michaelis-Menten kinetics of *Pb*-M1, compared to *Pf*A-M1 and *Pv*-M1 (inset table). *Pb*-M1 catalyzes Leu-Mec with higher affinity than *Pf*A-M1 and *Pv*-M1. **(F)** Michaelis-Menten kinetics of *Pb*-M17, compared to *Pf*A-M17 and *Pb*-M17. Affinity for Leu-Mec is conserved across the three homologs.

We confirmed *Pb*-M1 was active in the absence of supplemental metal ions (Fig 1C), but that *Pb*-M17 required supplementation with Co^2+^ or Mn^2+^ for optimal activity *in vitro* (Fig 1D). This fits with behavior observed from other M1 and M17 aminopeptidases, including *Pf*A- and *Pv*-M1 and M17. Although *Pb*-M17 demonstrated optimal activity in 0.5 mM Co^2+^ and 0.25-1.0 mM Mn^2+^ (Fig 1D), kinetic parameters were determined in the presence of 1 mM Co^2+^ to align with previously published experiments (11). The kinetic parameters of *Pb*-M1 were determined in the absence of metal ions. The affinity of *Pb*-M1 for Leu-Mec is approximately 3.5-fold higher than both *Pf*A-M1 and *Pv*-M1 (Fig 1E), but *Pb*-M17 is similar to that of its *Plasmodium* homologs (Fig 1F).

### Comparing the substrate specificity at the P1 position across the Plasmodium M1 aminopeptidases

To profile the P1 substrate specificities of the enzymes, we used the same fluorogenic substrate library that we have used previously to characterize *Pf*A-M1 (13) This library contains 19 natural and 42 non-natural amino acids coupled to an ACC leaving group. To compare enzyme selectivity, the amount of fluorescent end-product generated from each substrate within the experimental time frame was measured. The substrate that produced the highest fluorescence levels within the designated time frame was assigned a value of 100% and used to calculate relative digestion levels of other substrates (Fig 2). The results show that *Pv*-M1 and *Pb*-M1 are both most selective the same natural P1 amino acids (Ala, Arg, Leu, Lys, Met, Phe, Trp, Tyr) but their affinity for each is quite different. *Pv*-M1 had a P1 preference of Met>Ala>Leu before the charged residues Lys> Arg>>Gln. The enzymes were least active in removal of the larger aromatic side chains (Phe = Trp = Tyr) (Fig 2A). Interestingly, the preference for hydrophobic residues is reversed in *Pb*-M1 that preferred Lys/Arg before the bulkier hydrophobics Met>Leu and then Ala (Fig 2B). Including previous *Pf*A-M1 data in the comparison (Fig 2C), we can see that *Pf*A-M1 is more similar to that of *Pv*-M1, sharing a preference for the hydrophobic sidechains. *Pf*A-M1 appears to have a slightly broader profile as it shows activity toward smaller residues like Ser, Gly, Val and Thr (Fig 2C). A heat map of the three profile highlights that all three homologs appear to have little to no activity on residues with polar and uncharged sidechains, and moderate preference for aromatic residues (Fig 2D).

**Figure 2.**
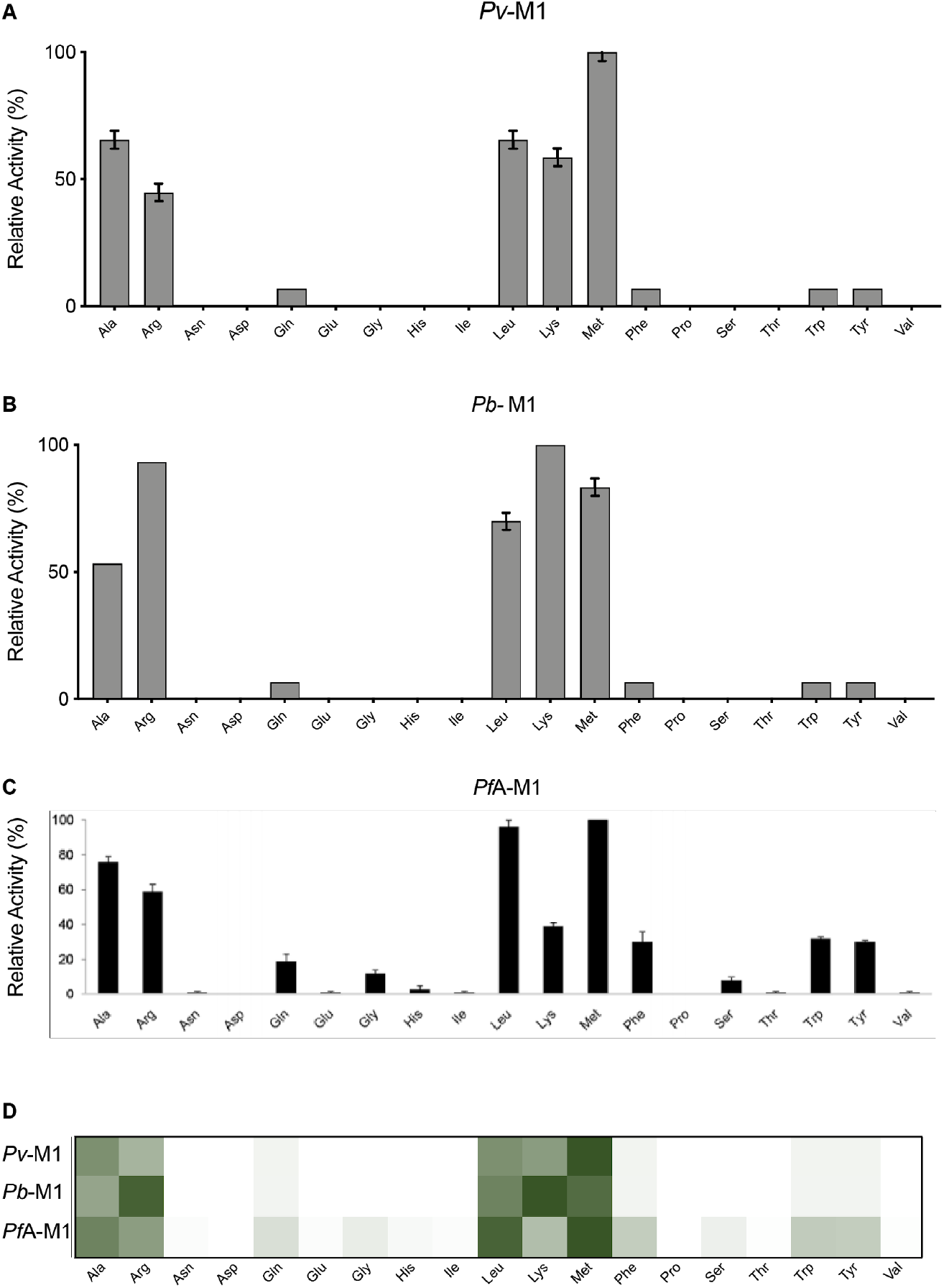
P1 Natural Amino Acid Substrate Profile of M1 Aminopeptidases from *P. falciparum, P. vivax* and *P. berghei*. P1 substrate specificity screening of *Pv*-M1 **(A)**, *Pb*-M1 **(B)** and *Pf*A-M1 **(C)** (15) with a 19-membered natural amino acid library. The amino acid with highest activity recorded was assigned as 100% activity, and activity against other amino acids is represented as percentage activity. **(D)** Heat map depicting the most favored amino acids as dark green and least favored as white, with all intermediate values depicted as shades of green.

When we investigated the extended library of 42 non-natural amino acids, we observed that *Pv*- and *Pb*-M1 have a very similar substrate specificity profile (Fig 3A, B). *Pv*-M1 had highest activity against hCha and hPhe, moderate activity against Nle and hTyr and low activity against hArg, Nva, styryl-Ala, allyl-Gly, 3-CN-Phe, Abu and hLeu. *Pb*-M1 shared this specificity, although had high activity against Arg derivative, hArg, and low activity against Ala derivative, Abu. The *Pb*-M1 preference for Arg over Ala is observed in both natural amino acids and their derivatives. The *Pf*A-M1 profile is quite different to both *Pv*-M1 and *Pb*-M1 (Fig 3C). At high activity levels, preferential cleavage of hCha, hArg and Nle is conserved, although at low activity levels *Pf*A-M1 exhibits a much broader specificity range. *Pf*A-M1 had low activity against substrates that are not hydrolyzed by *Pb*-M1 and *Pv*-M1, including Dap, cyclophentyl-Gly, (1-pyridin-4-yl)-Ala, Dab, 1-Nal, Phg, Bpa and neopentyl-Gly. Comparison of the heat map indicates that while *Pf*A-M1 has unique low-level activity against some non-natural residues, specificity is mostly conserved between the three homologs (Fig 3D).

**Figure 3.**
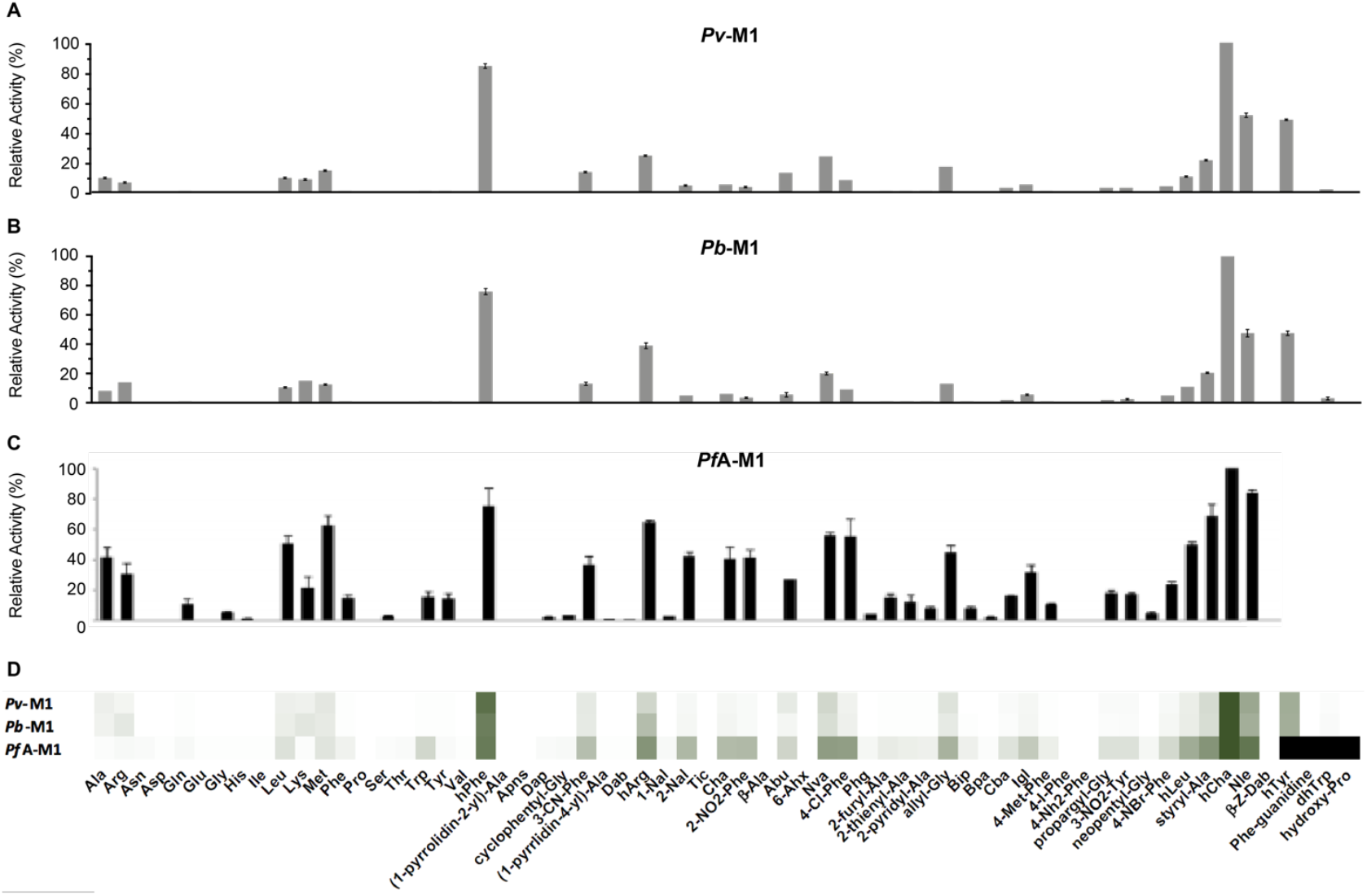
P1 Non-Natural Amino Acid Substrate Profile of M1 Aminopeptidases from *P. falciparum, P. vivax* and *P. berghei*. P1 substrate specificity screening of *Pv*-M1 **(A),** *Pb*-M1 **(B)** and *Pf*A-M1 **(C)** (15) 42-membered non-natural amino acid library. The amino acid with highest activity recorded was assigned as 100% activity, and activity against other amino acids is represented as percentage activity. **(D)** Heat map depicting the most favored amino acids as dark green and least favored as white, with all intermediate values depicted as shades of green.

The kinetic parameters of *Pv*-M1 and *Pb*-M1 were evaluated using the most preferred substrate, hCha (Table 1). *Pv*-M1 and *Pb*-M1 affinities and substrate turnover values were within 2-fold, although *Pv*-M1 had slightly higher affinity and turnover rates (Table 1). As a result, *Pv*-M1 had a marginally higher catalytic efficiency than *Pb*-M1 (*Pv*-M1 *k*_cat_ / *K*_m_ = 7.6 x 10^5^ M^−1^s^−1^; *Pb*-M1 *k*_cat_ / *K*_m_ = 3.0 x 10^5^ M^−1^s^−1^). Previously determined data shows that *Pf*A-M1 processes the same substrate with 15 to 40-fold less efficiency (*Pf*A-M1 *k*cat / *K*_m_ = 1.7 x 10^4^ M^−1^s^−1^), largely due to a 10 to 14-fold decrease in substrate turnover rate.

**Table 1.** Kinetic parameters of *Pf*A-M1, *Pb*-M1 and *Pv*-M1 against non-natural amino acid substrate, hCha. Assays carried out in biological triplicate and reported as mean ± standard error of the mean (SEM).

|  | *PfA*-M1 |  |  | *Pv*-M1 |  |  | *Pb*-M1 |  |  |
| --- | --- | --- | --- | --- | --- | --- | --- | --- | --- |
| | $K_m$<br>($\mu\text{M}$) | $k_{cat}$<br>($\text{s}^{-1}$) | $k_{cat}/K_m$<br>($\text{M}^{-1}\cdot\text{s}^{-1}$) | $K_m$<br>($\mu\text{M}$) | $k_{cat}$<br>($\text{s}^{-1}$) | $k_{cat}/K_m$<br>($\text{M}^{-1}\cdot\text{s}^{-1}$) | $K_m$<br>($\mu\text{M}$) | $k_{cat}$<br>($\text{s}^{-1}$) | $k_{cat}/K_m$<br>($\text{M}^{-1}\cdot\text{s}^{-1}$) |
| hCha | 96.3 ± 16.2 | 1.6 ± 0.2 | 1.7 x 10<sup>4</sup> | 29.6 ± 4.4 | 22.4 ± 0.9 | 7.6 x 10<sup>5</sup> | 53.9 ± 1.5 | 16.4 ± 2.9 | 3.0 x 10<sup>5</sup> |

± SEM

### Probing the substrate length and processivity of the Plasmodium M1 Aminopeptidases

*Pf*A-M1 has been shown to process dipeptide and tripeptide substrates with varying sequences (14) and in a processive manner (15). To probe the ability of *Pv*-M1 and *Pb*-M1 to also act in a processive manner, we designed two fluorescently labelled substrates longer than the commercially available Leu-Mec; a leucine di-peptide ([Leu]_2_) and a tetra-peptide ([Leu]_4_) coupled to a fluorescent reporter (NHMec). The kinetic parameters of the M1 aminopeptidases were evaluated with both peptide substrates (in addition to the commercially available Leu-NHMec substrate) and compared to evaluate the catalytic response to increasing substrate length (Table 2). Leucine repeats were selected for these substrates as P1 leucine was favored by all three M1 aminopeptidases, as well as allowing the same peptides to be evaluated against the M17 aminopeptidases as discussed later in this paper.

**Table 2.**
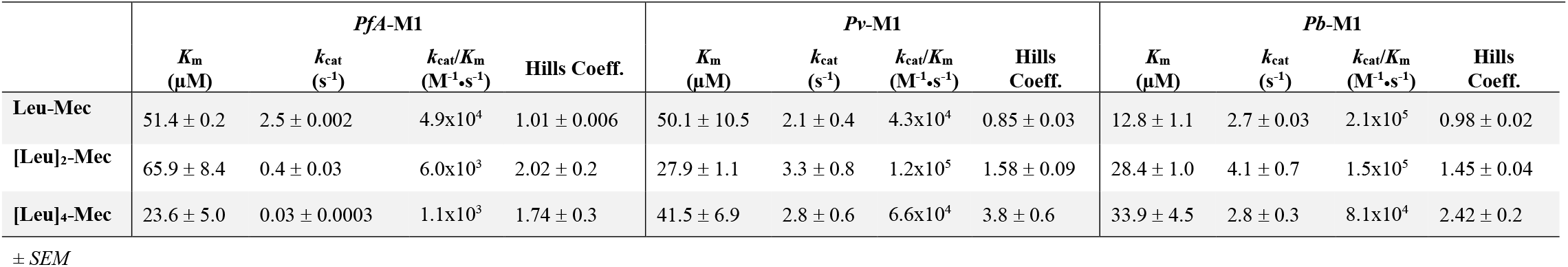
Kinetic parameters of *Pf*A-M1, *Pv*-M1 and *Pb*-M1 aminopeptidases against Leu-Mec, [Leu]_2_-Mec and [Leu]_4_-Mec. Assays carried out in biological triplicate and reported as mean ± standard error of the mean (SEM).

|  | *PfA*-M1 |  |  |  | *Pv*-M1 |  |  |  | *Pb*-M1 |  |  |  |
| --- | --- | --- | --- | --- | --- | --- | --- | --- | --- | --- | --- | --- |
| | $K_m$<br>($\mu\text{M}$) | $k_{cat}$<br>($\text{s}^{-1}$) | $k_{cat}/K_m$<br>($\text{M}^{-1}\cdot\text{s}^{-1}$) | Hills Coeff. | $K_m$<br>($\mu\text{M}$) | $k_{cat}$<br>($\text{s}^{-1}$) | $k_{cat}/K_m$<br>($\text{M}^{-1}\cdot\text{s}^{-1}$) | Hills Coeff. | $K_m$<br>($\mu\text{M}$) | $k_{cat}$<br>($\text{s}^{-1}$) | $k_{cat}/K_m$<br>($\text{M}^{-1}\cdot\text{s}^{-1}$) | Hills Coeff. |
| Leu-Mec | 51.4 ± 0.2 | 2.5 ± 0.002 | 4.9x10<sup>4</sup> | 1.01 ± 0.006 | 50.1 ± 10.5 | 2.1 ± 0.4 | 4.3x10<sup>4</sup> | 0.85 ± 0.03 | 12.8 ± 1.1 | 2.7 ± 0.03 | 2.1x10<sup>5</sup> | 0.98 ± 0.02 |
| [Leu]<sub>2</sub>-Mec | 65.9 ± 8.4 | 0.4 ± 0.03 | 6.0x10<sup>3</sup> | 2.02 ± 0.2 | 27.9 ± 1.1 | 3.3 ± 0.8 | 1.2x10<sup>5</sup> | 1.58 ± 0.09 | 28.4 ± 1.0 | 4.1 ± 0.7 | 1.5x10<sup>5</sup> | 1.45 ± 0.04 |
| [Leu]<sub>4</sub>-Mec | 23.6 ± 5.0 | 0.03 ± 0.0003 | 1.1x10<sup>3</sup> | 1.74 ± 0.3 | 41.5 ± 6.9 | 2.8 ± 0.6 | 6.6x10<sup>4</sup> | 3.8 ± 0.6 | 33.9 ± 4.5 | 2.8 ± 0.3 | 8.1x10<sup>4</sup> | 2.42 ± 0.2 |

± SEM

The M1 aminopeptidase homologs all showed an ability to processively cleave both the dipeptide and tetrapeptide, although each had unique kinetic responses to the increase in peptide substrate length. Affinity of *Pf*A-M1 for peptide substrates was not overly affected by substrate length, with all *K*_m_ values being within 2 to 3-fold of each other. *Pf*A-M1 turnover rate decreased ~100-fold as peptide length increased from one to four residues, slowing from a turnover rate of 2.5 s^−1^ for Leu-Mec to 0.03 s^−1^ for [Leu]_4_-Mec (Table 2). The steady decrease in turnover rate as substrate length increased means the catalytic efficiency also decreased by ~50-fold (Table 2). The *Pv*-M1 substrate selectivity profile also showed no relationship between increasing substrate length and affinity, and again all *K*_m_ values were within 2-fold of each other; Leu-Mec = 50.1 μM; [Leu]_2_-Mec = 27.9 μM; [Leu]_4_-Mec = 41.5 μM (Table 2). Changes in substrate length had little effect on substrate turnover and all three substrate *kcat* values were comparable; [Leu]_2_-Mec = 3.3 s^−1^, Leu-Mec = 2.1 s^−1^ and [Leu]_4_-Mec = 2.8 s^−1^ (Table 2). *Pb*-M1 was the only homolog where substrate length and affinity exhibited a clear correlation, with affinity steadily decreasing from 12.8 μM to 28.4 μM to 33.9 μM as substrate length increased (Table 2). *Pb*-M1 showed a similar turnover rate trend to *Pv*-M1, hydrolyzing [Leu]_2_-Mec the fastest (4.1 s^−1^), followed by [Leu]_4_-Mec (2.8 s^−1^) and Leu-Mec (2.7 s^−1^). *Pb*-M1 became less catalytically efficient as substrate length increased, owing to the steady decrease in substrate affinity.

*Pf*A-M1 activity is most commonly characterized using the standard Leu-Mec substrate and has never been shown to act cooperatively to process substrates (Hill’s coefficient (h) = 1.01, Table 2). However, we observed that when substrate length increased to two residues, the Hill’s coefficient (h) similarly increased to h = 2.02, indicating that *Pf*A-M1 processed [Leu]_2_-Mec cooperatively (Table 2). The Hill’s coefficient dropped with the increase to four residues (h = 1.74), meaning *Pf*A-M1 was still acting cooperatively, but maybe to a lesser degree (Table 2). *Pv*-M1 and *Pb*-M1 both functioned non-cooperatively against Leu-Mec, with Hill’s coefficients of 0.85 and 0.98 respectively (Table 2). Similar to *Pf*A-M1, both *Pv*-M1 and *Pb*-M1 aminopeptidases became cooperative when substrate length increased to two residues (*Pv*-M1 h = 1.58, *Pb*-M1 h = 1.45, Table 2), however, continued to become more cooperative as substrate length increased to four residues (*Pv*-M1 h = 3.8, *Pb*-M1 h = 2.42, Table 2).

### M1 aminopeptidases demonstrate intra-peptide sequence specificity

The fact that our M1 aminopeptidases were able to cleave [Leu]_4_-Mec was unsurprising as previous studies had shown that *PfA-M1* could degrade hexapeptides derived from hemoglobin (a chain peptide VDPENF and ß chain peptide VDPVNF) (Harbut et al., 2011). Using this information, we produced LDPENF as a derivative of the a chain natural substrate and assessed the activity of the three M1 aminopeptidases toward this peptide (Fig 4). Using mass spectrometry to detect the end point product of digestion, we were able to show that all three *Plasmodium* M1 enzymes were able to cleave the P1 Leu from this hexapeptide (Fig 4). We also anticipated that the P1’ Asp residue in our peptide sequence would prevent processive activity, as none of our three enzymes were able to efficiently digest a P1 Asp or P1’ Pro. This was indeed the case, as DPENF was the only smaller peptide fragment detected from our end point experiments (Supp Fig S1).

**Figure 4.**
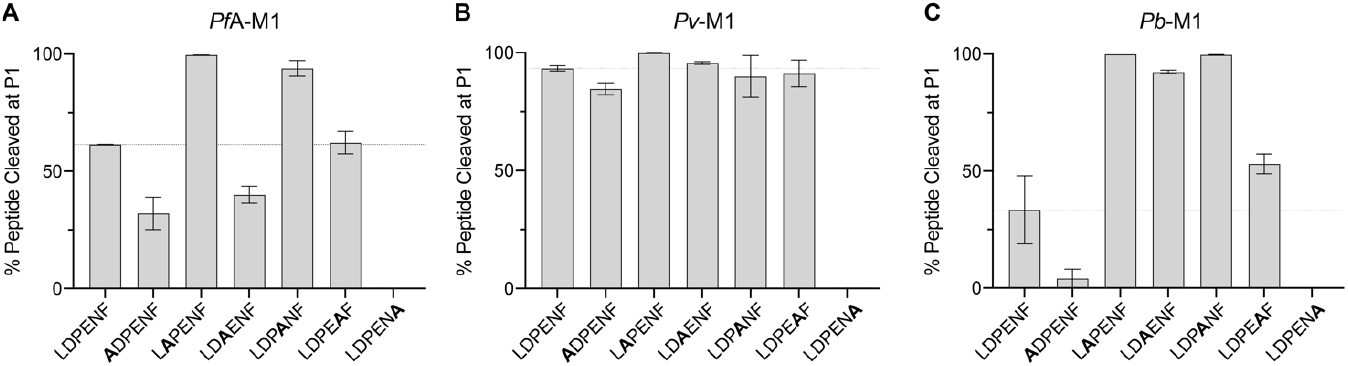
Activity of *Pf*A-M1, *PV*-M1 and *Pb*-M1 against hemoglobin-derived hexapeptide and alanine screen peptides. Activity of *Pf*A-M1 (A), *Pv*-M1 (B) and *Pb*-M1 (C) against hemoglobin derived hexapeptide, LDPENF, and alanine screen peptides, ADPENF, LAPENF, LDAENF, LDPANF, LDPEAF and LDPENA. Activity presented as percentage of peptide cleaved at the P1 position.

**Figure 5.**
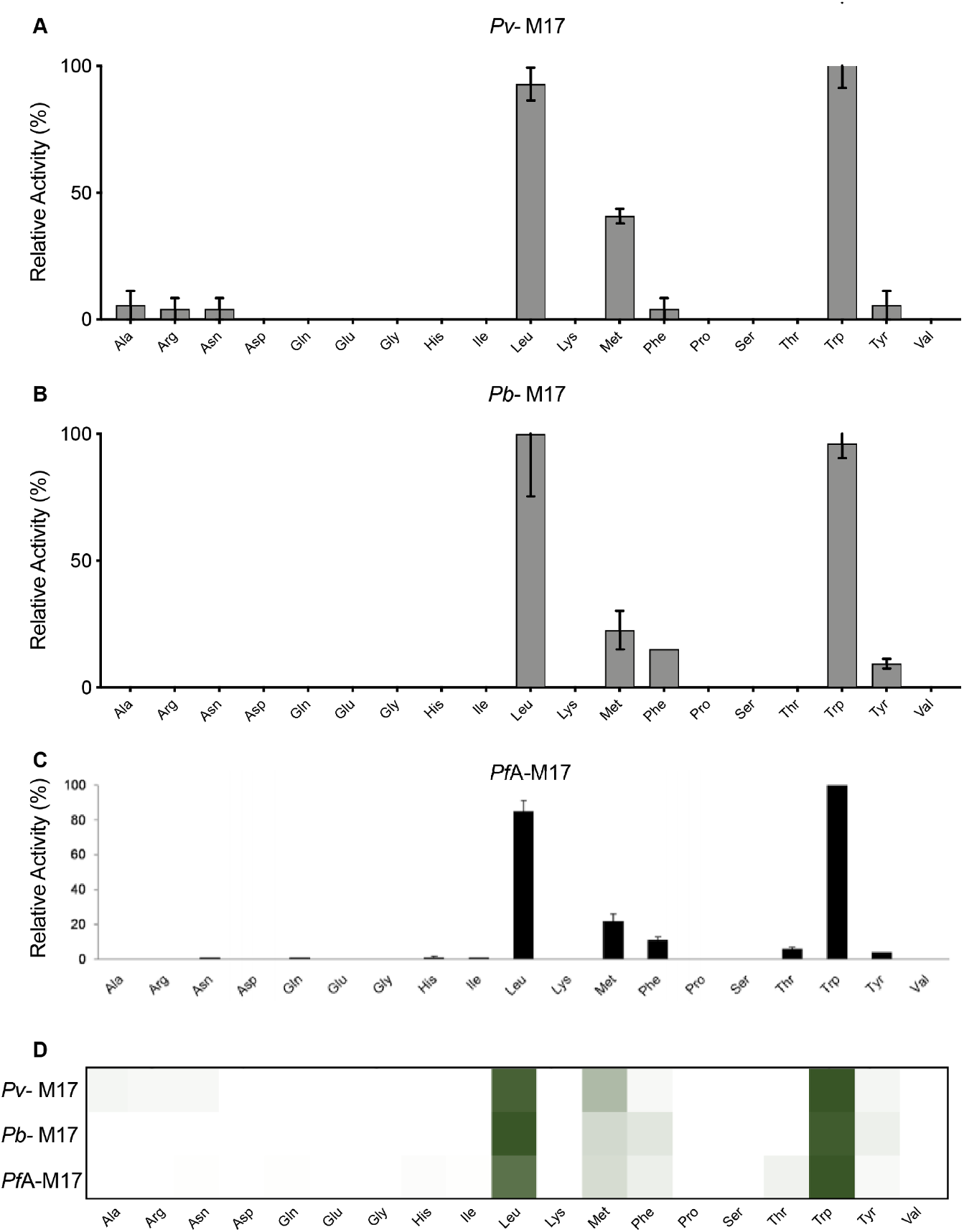
P1 Natural Amino Acid Substrate Profile of M17 Aminopeptidases from *P. falciparum, P. vivax* and *P. berghei*. P1 substrate specificity screening of *Pv*-M17 **(A)**, *Pb*-M17 **(B)** and *Pf*A-M17 **(C)** (15) with a 19-membered natural amino acid library. The amino acid with highest activity recorded was assigned as 100% activity, and activity against other amino acids is represented as percentage activity. **(D)** Heat map depicting the most favored amino acids as dark green and least favored as white, with all intermediate values depicted as shades of green.

Florent (2001) and Dalal (2012) both demonstrated that *Pf*A-M1 has specificity beyond the P1 site. The digestible LDPENF hexapeptide was used as a template for a peptide library designed to test the intra-peptide specificities of *Pf*A-M1, *Pv*-M1 and *Pb*-M1. An alanine screen library was produced, with each peptide substituting Ala at a different position to gain an understanding of the accepted residues at each position. Peptide cleavage was detected using mass spectrometry and the percentage cleavage of each peptide calculated. Assay times were reduced for *Pv*-M1 and *Pb*-M1 as all peptides were cleaved to almost 100% within the 45 min timeframe, resulting in a loss of specificity information (Supp Fig S2).

*Pf*A-M1 showed a preference for LAPENF and LDPANF over LDPENF. When Ala replaced Asp in the P1’ position, activity increased from 61% to 100%. Similarly, the substitution of Glu for Ala in the P3’ position resulted in increased activity (94%). The peptide product, APENF, produced from the cleavage of LAPENF was not further digested due to the inability of *Pf*A-M1 to cleave peptide with a P1’ Pro. *Pf*A-M1 appears to function more efficiently when negatively charged residues are replaced, regardless of their position within the peptide. Replacement of P1 Leu with Ala resulted in a decrease in activity from 61% to 32%, as *Pf*A-M1 prefers Leu in the P1 position (a result supported by the P1 screen). Substitution of the P2’ Pro residues similarly resulted in decreased activity (40%), suggesting that the rigidity Pro residue provides to the peptide may be vital for optimal activity. LDPENA was the only peptide that *Pf*A-M1 did not cleave, which is likely a result of replacing a large aromatic residue with the comparatively small Ala sidechain. *Pv*-M1 was incredibly efficient at cleaving the hemoglobin-derived peptide and alanine screen peptides. Even with the reduction of enzyme concentration, peptide concentration and incubation time, activity rates were in excess of 80% for each peptide, except for LDPENA, which was not processed at all. *Pv*-M1 demonstrated a lower tolerance for Ala in the P1 position than Leu, with activity rates dropping from 93% to 84%. The substitution of Ala in the P1’ position in place of Asp to produce LAPENF increased activity to 99%, while the activity levels of LDAENF, LDPANF and LDPEAF remained approximately similar to LDPENF. *Pb-M1* preferentially cleaved both LAPENF and LDPANF over LDPENF. *Pb*-M1 had 99% activity against both LAPENF and LDPANF, compared to the 33% activity against LDPENF, thereby exhibiting a preference for peptide with fewer negatively charged residues. *Pb*-M1 also showed a preference for Ala in the P2’ position in place of Pro, the substitution increasing activity to 92%, indicating that increased peptide flexibility was advantageous for enzyme activity. *Pb*-M1 showed decreased activity against both ADPENF and LDPENA. *Pb*-M1 catalyzed ADPENF with activity rate of only 4%, while there was no observed cleavage of LDPENA.

### The Plasmodium M17 aminopeptidase P1 profile is highly conserved

The P1 substrate specificity of the M17 aminopeptidases was profiled using the same 19-membered natural amino acid and 42-membered non-natural amino acid library used to profile the M1 aminopeptidases. Compared to the M1 homologs, the M17 aminopeptidases had a very limited substrate specificity range. *Pv*-M17 demonstrated a preference for the bulky nonpolar residues Trp, Leu and Met and showed low-level activity (<10%) against Ala, Arg, Asn, Phe and Tyr (Fig 6A). *Pb*-M17 demonstrated a similar preference for the bulky hydrophobic residues, although catalyzed the removal of Leu with highest efficiency followed by Trp and Met. *Pb*-M17 was less efficient at processing Met than *Pv*-M17 was but showed higher activity against aromatic residues Phe and Tyr (Fig 6B). When comparing with the previously determined *Pf*A-M17 data, we can see the profiles are very much conserved between homologs (Fig 6C). The heat map indicated a conserved preference for Leu and Trp, followed by Met, and finally Phe and Tyr (Fig 6D). *Pv*-M17 and *Pf*A-M17 both had low-level activities against several substrates that was not conserved between homologs.

**Figure 6.**
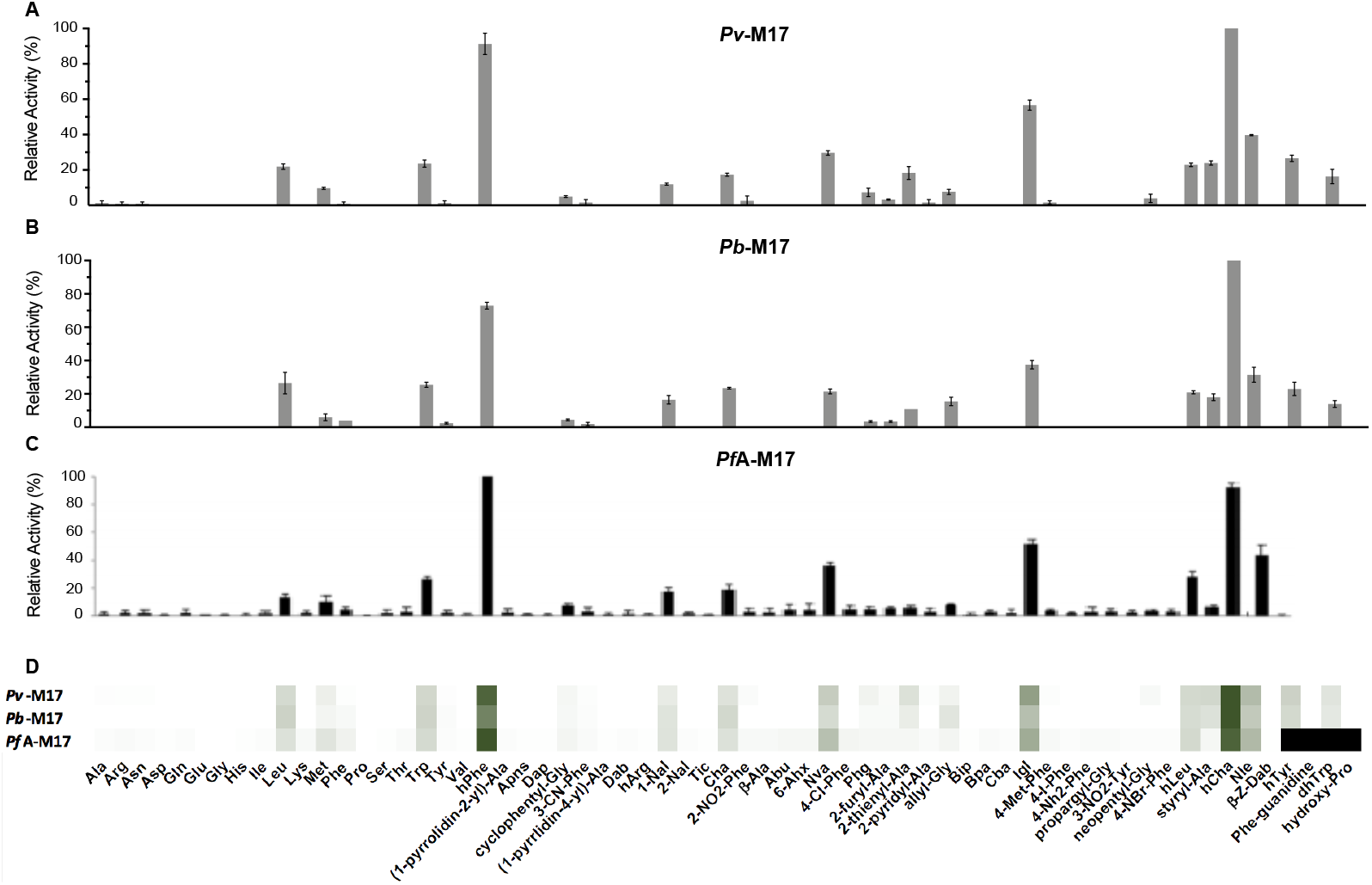
P1 Non-Natural Amino Acid Substrate Profile of M17 Aminopeptidases from *P. falciparum, P. vivax* and *P. berghei*. P1 substrate specificity screening of *Pv*-M17 **(A),** *Pb*-M17 **(B)** and *Pf*A-M17 **(C)** (15) 42-membered non-natural amino acid library. The amino acid with highest activity recorded was assigned as 100% activity, and activity against other amino acids is represented as percentage activity. **(D)** Heat map depicting the most favored amino acids as dark green and least favored as white, with all intermediate values depicted as shades of green.

The M17 aminopeptidases cleaved the non-natural amino acids with much higher efficiency than the natural amino acids and activity profiles were again largely conserved across the three homologs. *Pv*-M17 and *Pb*-M17 demonstrated similar specificity profiles; both aminopeptidases preferentially cleaved hCha > hPhe > Igl > Nle with highest efficiency, and cleaved hCha with 4-fold higher efficiency than the best natural amino acid (Fig 7A, 7B). *Pv*-M17 then preferentially cleaved Nva > hTyr > styryl-Ala > hLeu, while *Pb*-M17 preferentially cleaved Cha > hTyr > Nva > hLeu. *Pv*-M17 had higher activity against Val and Ala derivatives Nva and styryl-Ala (Fig 7A), while *Pb*-M17 had a unique preference for Ala derivative Cha (Fig 7B). When comparing to the previously determined *Pf*A-M17 data, the strong preferences observed for hPhe, hCha, Igl and Nle are conserved across the three homologs. *Pf*A-M17 also showed higher activity levels against the non-natural amino acids compared to the natural amino acids, cleaving the most preferred non-natural substrate (hPhe) with 4-fold higher efficiency than the best natural substrate (Trp). The heat map indicates that the substrates processed with high efficiency were conserved across the three homologs, whereas the substrates that are processed with low efficiency are more unique to each homolog (Fig 8D).

**Figure 7.**
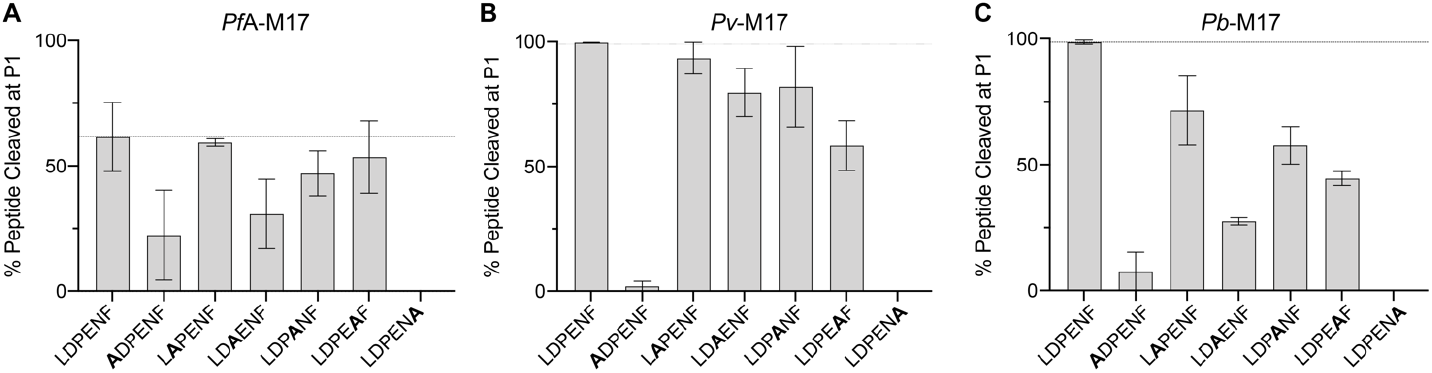
Activity of *Pf*A-M17, *Pv*-M17 and *Pb*-M17 against hemoglobin-derived hexapeptide and alanine screen peptides. Activity of *Pf*A-M17 (A), *Pv*-M17 (B) and *Pb*-M17 (C) against hemoglobin derived hexapeptide, LDPENF, and alanine screen peptides, ADPENF, LAPENF, LDAENF, LDPANF, LDPEAF and LDPENA. Activity presented as percentage of peptide cleaved at the P1 position.

**Figure 8.**
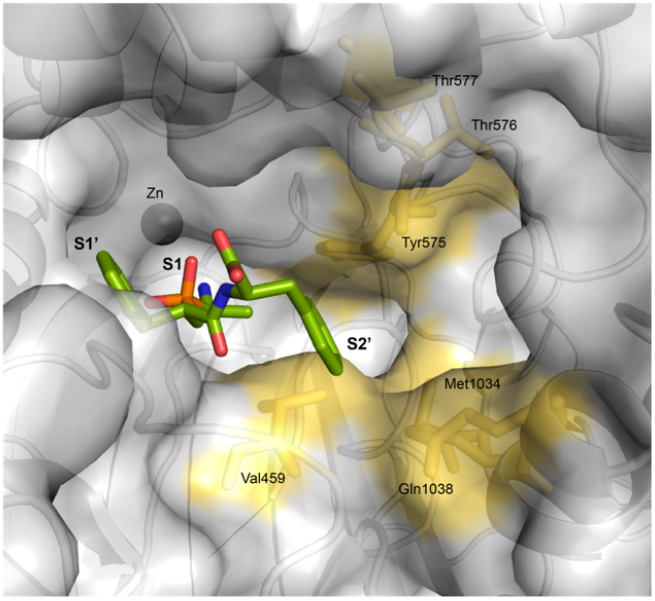
APN inhibitor PL250 docked into *PfA-M1* X-ray crystal structure with P2’ subsite highlighted. *E. coli* APN X-ray crystal structure in complex with PL250 (PDB ID: 2ZXG) used to dock PL250 (green sticks) into *Pf*A-M1 X-ray crystal structure (PDB ID: 3EBG). S1, S1’ and S2’ pockets labelled. Residues postulated to contribute to the *Pf*A-M1 S2’ pocket are shown as yellow sticks and surface colored yellow.

For each of the homologs, we determined the kinetic parameters for the non-natural amino acid substrate, hCha (Table 3). *Pv*-M17 and *Pb*-M17 had remarkably similar kinetic parameters, with affinities of 1.1 μM and 1.2 μM respectively and turnover rates of 0.4 and 0.3 s^−1^ respectively. Together, these result in catalytic efficiencies of 3.6 x 10^5^ and 2.5 x 10^5^ M^−1^s^−1^ respectively. The previously determined *Pf*A-M17 results illustrate a slightly higher catalytic efficiency (4.8 x 10^5^ M^−1^ s^−1^) for hCha than *Pv*-M17 and *Pv*-M17, which was largely due to the higher affinity of *Pf*A-M17 for hCha; *K*_m_ = 0.44 μM (Table 3).

**Table 3.** Kinetic parameters of *Pf*A-M17, *Pb*-M17 and *Pv*-M17 against unnatural amino acid substrate hCha. Assays carried out in biological triplicate and reported as mean *±* standard error of the mean (SEM).

|  | *PfA*-M17 |  |  | *Pv*-M17 |  |  | *Pb*-M17 |  |  |
| --- | --- | --- | --- | --- | --- | --- | --- | --- | --- |
| | $K_m$<br>($\mu\text{M}$) | $k_{cat}$<br>($\text{s}^{-1}$) | $k_{cat}/K_m$<br>($\text{M}^{-1}\cdot\text{s}^{-1}$) | $K_m$<br>($\mu\text{M}$) | $k_{cat}$<br>($\text{s}^{-1}$) | $k_{cat}/K_m$<br>($\text{M}^{-1}\cdot\text{s}^{-1}$) | $K_m$<br>($\mu\text{M}$) | $k_{cat}$<br>($\text{s}^{-1}$) | $k_{cat}/K_m$<br>($\text{M}^{-1}\cdot\text{s}^{-1}$) |
| hCha | $0.44 \pm 0.02$ | $0.21 \pm 0.005$ | $4.8 \times 10^5$ | $1.1 \pm 0.03$ | $0.4 \pm 0.04$ | $3.6 \times 10^5$ | $1.2 \pm 0.1$ | $0.3 \pm 0.03$ | $2.5 \times 10^5$ |

± SEM

### Pv-M17 and Pb-M17 process dipeptides and tripeptides with greater efficiency than PfA-M17

The same Leu-based peptides used to characterise M1 processivity and kinetic activity were used to analyse the kinetic behaviours of the M17 homologs. Each of the M17 aminopeptidases successfully cleaved the Leu-based dipeptide and tetrapeptides processively, with kinetic responses to increased substrate length highly conserved compared to the M1 homologs. *Pf*A-M17 had the highest affinity for [Leu]_4_-Mec (*K*_m_ = 6.9 μM), although processed this substrate with the slowest turnover rate (0.02 s^−1^). [Leu]_2_-Mec was processed by *Pf*A-M17 with the lowest affinity (*K*_m_ = 30.8 μM) but the highest turnover rate (0.1 s^−1^), indicating an inverse relationship between substrate affinity and turnover rate (Table 4). As substrate length increased, *Pf*A-M17 catalytic efficiency progressively decreased and Leu-Mec was catalyzed (4.6 x 10^3^) with ~2-fold higher efficiency than [Leu]_4_-Mec (2.6 x 10^3^) (Table 4). *Pv*-M17 catalyzed Leu-Mec with both the highest affinity (20.1 μM) and highest turnover rate (0.9 s^−1^). [Leu]_4_-Mec was catalyzed with the same affinity (22.6 μM), but the slowest turnover rate (0.4 s^−1^), and [Leu]_2_-Mec was catalyzed with the lowest affinity (32.9 μM) and the same turnover rate to that of Leu-Mec (0.8 s^−1^). As seen with *Pf*A-M17, *Pv*-M17 decreased catalytic turnover as substrate length increased, catalyzing Leu-Mec with ~3-fold higher efficiency than [Leu]_4_-Mec (Table 4). *Pb*-M17 had similar affinity behaviors to *Pf*A-M17, showing highest affinity for [Leu]_4_-Mec (4.6 μM) *versus* either Leu-Mec (19.4 μM) or [Leu]_2_-Mec (25 μM). *Pb*-M17 turnover rate slowed dramatically as peptide length increased, with a ~23-fold decrease observed between Leu-Mec (11.4 s^−1^) and [Leu]_2_-Mec (0.5 s^−1^), and a further 6-fold decrease between [Leu]_2_-Mec and [Leu]_4_-Mec (0.08 s^−1^). The marked drop in turnover rates was reflected in catalytic efficiency, with Leu-Mec being processed with 31-fold more efficiency than [Leu]_4_-Mec.

**Table 4.**
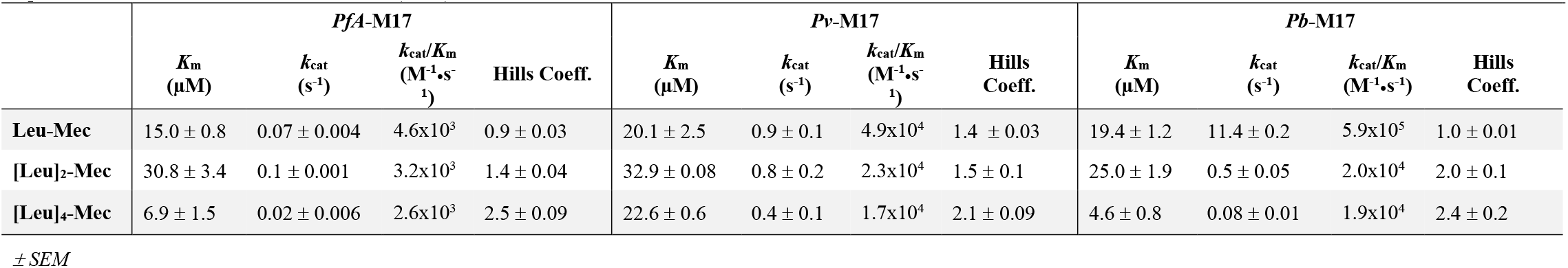
Kinetic parameters of *pf*A-M17, *Pv*-M17 and *Pb*-M17 aminopeptidases against Leu-Mec, [Leu]_2_-Mec and [Leu]_4_-Mec. Assays carried out in biological triplicate and reported as mean ± standard error of the mean (SEM).

|  | *PfA*-M17 |  |  |  | *Pv*-M17 |  |  |  | *Pb*-M17 |  |  |  |
| --- | --- | --- | --- | --- | --- | --- | --- | --- | --- | --- | --- | --- |
| | $K_m$<br>($\mu\text{M}$) | $k_{cat}$<br>($\text{s}^{-1}$) | $k_{cat}/K_m$<br>($\text{M}^{-1}\cdot\text{s}^{-1}$) | Hills<br>Coeff. | $K_m$<br>($\mu\text{M}$) | $k_{cat}$<br>($\text{s}^{-1}$) | $k_{cat}/K_m$<br>($\text{M}^{-1}\cdot\text{s}^{-1}$) | Hills<br>Coeff. | $K_m$<br>($\mu\text{M}$) | $k_{cat}$<br>($\text{s}^{-1}$) | $k_{cat}/K_m$<br>($\text{M}^{-1}\cdot\text{s}^{-1}$) | Hills<br>Coeff. |
| Leu-Mec | $15.0 \pm 0.8$ | $0.07 \pm 0.004$ | $4.6 \times 10^3$ | $0.9 \pm 0.03$ | $20.1 \pm 2.5$ | $0.9 \pm 0.1$ | $4.9 \times 10^4$ | $1.4 \pm 0.03$ | $19.4 \pm 1.2$ | $11.4 \pm 0.2$ | $5.9 \times 10^5$ | $1.0 \pm 0.01$ |
| [Leu]<sub>2</sub>-Mec | $30.8 \pm 3.4$ | $0.1 \pm 0.001$ | $3.2 \times 10^3$ | $1.4 \pm 0.04$ | $32.9 \pm 0.08$ | $0.8 \pm 0.2$ | $2.3 \times 10^4$ | $1.5 \pm 0.1$ | $25.0 \pm 1.9$ | $0.5 \pm 0.05$ | $2.0 \times 10^4$ | $2.0 \pm 0.1$ |
| [Leu]<sub>4</sub>-Mec | $6.9 \pm 1.5$ | $0.02 \pm 0.006$ | $2.6 \times 10^3$ | $2.5 \pm 0.09$ | $22.6 \pm 0.6$ | $0.4 \pm 0.1$ | $1.7 \times 10^4$ | $2.1 \pm 0.09$ | $4.6 \pm 0.8$ | $0.08 \pm 0.01$ | $1.9 \times 10^4$ | $2.4 \pm 0.2$ |

± SEM

Similar to M1, the kinetic behaviors of *Pf*A-M17 have been routinely characterized using single residues linked to a fluorescent reporter, i.e. Leu-Mec. *Pf*A-M17 has never been reported to behave cooperatively when using these single residue substrates, and we similarly saw no evidence of cooperativity when using Leu-Mec (Table 4). However, when using [Leu]_2_-Mec and [Leu]_4_-Mec, we observed Hill’s coefficients of 2.02 and 1.74 respectively, showing for the first time that *Pf*A-M17 functions cooperatively in the presence of longer peptide substrates. *Pv*-M17 and *Pb*-M17 showed very similar cooperativity behaviors; each processed Leu-Mec non-cooperatively with Hill’s coefficients of 0.85 and 0.98 respectively. As substrate length increased to a dipeptide, the *Pv*-M17 and *Pb*-M17 Hill’s coefficients similarly increased to 3.29 and 2.5 respectively, and upon increasing to a tetrapeptide the Hill’s coefficients increased again to 3.8 and 4.6 respectively. The continual increase of Hill’s coefficients indicates that the aminopeptidases operate in an increasingly cooperative manner as substrate length increases.

### Intrapeptide sequence specificity is highly conserved between M17 homologs, and integral to catalytic efficiency

We used the same Ala peptide library screen to assess intrapeptide specificity within the M17 aminopeptidases. Similar to the M1s, *Pf*A-M17 digested peptides at a much slower rate compared to *Pv*-M17 and *Pb*-M17. To account for this, *Pf*A-M17 experiments were carried out for a total of 45 mins, whereas *Pv*-M17 and *Pb*-M17 experiments were carried out for 10 mins (*Pv*-M17 and *Pb*-M17 digested all peptides to almost 100% in the 45 min time frame, Supp Fig S3). All experiments were carried out in technical duplicate; however, we did observe significant error in the two *Pf*A-M17 replicates. Despite the large degree of error, the two replicates had a conserved order of preference for the peptides tested. (LAPENF/LDPENF/LDPEAF > LDPANF > LDAENF > ADPENF > LDPENA) (Fig 7A). *Pf*A-M17 cleaved the hemoglobin-derived hexapeptide with 62% efficiency and did not show enhanced activity for any other peptides. LAPENF, LDPANF and LDPEAF were cleaved with similar efficiency to LDPENF, at rates of 59%, 47% and 53% respectively. The only substitutions that resulted in a change in activity were at the P1, P2’ and P5’ positions (Fig 7A). The substitution of Leu for Ala in the P1 position is clearly unfavorable, with activity rates dropping to 22% due to the pre-determined preference for Leu in the P1 position over Ala. Replacing the P2’ Pro with Ala reduced activity to 31%, likely due to the increase in peptide flexibility that an Ala in that position provides. The substitution of Phe for Ala in the P5’ position had the most dramatic effect on substrate specificity, reducing activity to <1%. Like *Pf*A-M1, *Pf*A-M17 appears to require a bulky aromatic residue on the C-terminal to successfully cleave the peptide substrate. *Pv*-M17 cleaved LDPENF with the highest efficiency of all peptides, and any substitutions to the sequence resulted in no change in activity or in some cases, a decrease in activity (Fig 7B). LAPENF and LDPANF were within a similar activity rate range to LDPENF, indicating that P1’ and P3’ positions tolerate Ala the same as, or slightly less than, Asp and Glu respectively. *Pv*-M17 had a unique tolerance for Ala in the P2’ position in place of Pro, with activity rates of 80%. Substitution of Asn to Ala in the P4’ position reduced activity by almost half from 99% to 58%, while substitution to Ala in the P1 and P5’ essentially ablated all activity. *Pb*-M17 also digested LDPENF with the highest efficiency with 98% cleaved, however showed a decrease in activity against LAPENF and LDPANF that was unique amongst the three homologs (Fig 7C). LAPENF was digested to 71%, and LDPANF digested to 57%, compared to the 98% cleavage rate before substitution. Substitution of Pro and Asn at the P2’ and P4’ reduced activity to 24% and 44%, and substitution for Leu and Phe in the P1 and P5’ positions further reduced activity to 7% and 0%.

## Discussion

As drug resistance to frontline antimalarials continues to proliferate, there is an urgent need to generate new agents toward novel drug targets (19). *Plasmodium* M1 and M17 aminopeptidases are validated targets for the design of novel anti-malarial agents, and the subject of ongoing structure-based drug design programs (8–11, 20–23). The development of potent and selective inhibitors requires a thorough characterization and comparison of how the target aminopeptidases function. Profiling kinetic fingerprints and identifying divergent versus conserved features provides insight into the inhibition of enzymes across families are species. Furthermore, by profiling the *P. berghei* enzymes in addition to *P. falciparum* and *P. vivax,* we are ensuring that structure-activity relationship data is applicable to relevant animal (mouse) trials. The work we have presented here sheds light on kinetic characteristics that are conserved within the M1 and M17 aminopeptidase families that can be exploited to achieve cross-species inhibition.

M1 aminopeptidases possess a broad P1 specificity profile divided into two categories; favoring charged P1 residues, as seen with *Pb*-M1, (24, 25), or favoring hydrophobic P1 residues, as seen with *Pf*A-M1 and *Pv*-M1 (24, 26). P1 specificity was generally conserved between homologs, although *Pf*A-M1 demonstrated a broader acceptance of residues at the P1 position than either *Pv*-M1 and *Pb*-M1. This breadth of specificity, however, appeared to come at a cost to substrate turnover rate. Discrepancy in turnover rate between *Pf*A-M1 and *Pv*-M1/*Pb*-M1 was exacerbated as substrate length increased; Leu-Mec was processed similarly by the three substrates, however, *Pf*A-M1 processed [Leu]_2_-Mec at rates ~10-fold lower and [Leu]_4_-Mec at rates ~100-fold than *Pv*-M1 and *Pb*-M1. *Pf*A-M1 also required at least 4-fold longer incubation time to achieve similar levels of hexapeptide digestion. All residues that are involved in active site metal-coordination and architecture are conserved between *Pf*A-M1, *Pv*-M1 and *Pb*-M1, suggesting that differences in catalytic efficiency are unlikely a result of the hydrolysis reaction. Further, affinities for the various substrates assessed remained largely consistent amongst homologs, indicating that substrate binding is also conserved.

M1 family aminopeptidases undergo large-scale dynamic changes between an open conformation, which exposes the internal active site, and closed conformation (16, 27, 28). When in the open conformation, the active site is disordered and the enzymes are inactive, suggesting that the active site must bind peptide substrates to induce closing and recovery of catalytic activity (27, 28). Upon completion of the hydrolysis reaction, the opening motion mediates release of the hydrolyzed products and recruitment of new substrates. The speed at which the enzyme can process substrates therefore relies – to some degree – on how quickly the enzyme can transition between these open and closed states. *Pf*A-M1 exhibited decreased turnover rates when compared to *Pv*-M1 and *Pb*-M1, suggesting the transition between open and closed conformations may be slower, leaving the active site is exposed for an extended period of time. This could allow time for less favorable substrates to bind, resulting in a broad specificity profile. Similarly, we postulate that the faster substrate turnover rates of *Pv*-M1 and *Pb*-M1 correlates to an increase in dynamic behavior. However, rapid transition between conformations may limit the opportunity to bind less preferred substrates, resulting in a faster enzyme with a narrower substrate specificity.

In addition to conformational regulation of proteolysis, M1 aminopeptidases have been suggested to possess an allosteric site that is separate to but in close proximity to the active site (16, 29). Activation of the allosteric site is proposed to occur through a single long peptide substrate that reaches both the active site and allosteric site, as is the case for ERAP1, which shows length specificity for substrates long enough to reach both sites (9-16 residues) (16, 27, 29). Alternatively, activation is postulated to occur by simultaneous occupation of the active and allosteric sites by two small peptides. Our results show that as the leucine-based substrate length increased, cooperativity increased. We postulate that both the active site and a separate secondary site were simultaneously occupied by two individual peptides. Peptide binding in the secondary site did not overly improve catalytic efficiency, therefore is likely to have a function other than activating catalytic activity. Instead this secondary site may function as a dynamic regulation sight with peptide binding signaling the enzyme to close and initiate hydrolysis. Yeast LTA4H demonstrates this type of ‘induced fit’ mechanism as coordination of bestatin in the active site induces the closing motion and structural ordering of the active site (30). Cooperativity of each M1 homolog increased as peptide length increased, suggesting that longer peptides are increasingly efficient at accessing and occupying the secondary site. *Pf*A-M1, *Pv*-M1 and *Pb*-M1 each exhibited unique catalytic responses to the increase in peptide substrate length, meaning the extent or process by which the secondary site mediates dynamicity or induces structural change is likely specific to each enzyme.

*Pf*A-M1 has been shown to digest a hemoglobin hexa-peptide in parasites (4). The M1 homologs digested the hemoglobin-derived hexapeptide and all of the alanine screen peptides, with the exception of LDPENA. *Pv*-M1 and *Pb*-M1 continued to demonstrate elevated substrate turnover rates in comparison to *Pf*A-M1, with *Pv*-M1 exhibiting extremely fast turnover rates relative to the other two homologs. *Pv*-M1 showed the least amount of intrapeptide specificity, likely a byproduct of the rapid turnover rate. The M1 aminopeptidases demonstrated a conserved preference for Leu over Ala in the P1 position, a result that is supported by the P1 substrate screen. The M1 homologs shared a conserved preference for peptide substrates LAPENF and LDPANF.

The active site and substrate binding cleft of *Pf*A-M1 is highly hydrophobic and therefore has a low tolerance for negatively charged residues such as Asp and Glu. These residues were also unfavorable in the P1 position, as none of the homologs successfully processed the Asp or Glu substrates (except for *Pf*A-M1, which shows minimal activity against Glu) (Fig 2). The only difference observed between the three homologs was the digestion of LDAENF; *Pv*-M1 and *Pb*-M1 both favored the removal of the P2’ Pro, whereas the removal was detrimental to *Pf*A-M1 activity. We postulated that this may be due to structural differences in the S2’ binding sites of *Pv*-M1 and *Pb*-M1 compared to *Pf*A-M1 that would favor an Ala in place of a Pro. The S2’ binding site of *Pf*A-M1 has not previously been mapped. The *E. coli* APN crystal structure in complex with inhibitory molecule, PL250, has a defined S2’ site, and was superimposed onto the *Pf*A-M1 structure to align the S2’ site (14, 31). We estimated that the *Pf*A-M1 S2’ binding pocket is lined by residues Val459, Tyr575, Thr576 and Gln1038 (Fig 8). These residues are conserved in *Pv*-M1 and *Pb*-M1, indicating that their S2’ site structure is also likely conserved. The difference in activity against LDAENF may be due therefore to the difference in dynamic nature between the homologs, with *Pv*-M1 and *Pb*-M1 accepting flexible peptides with a higher tolerance than *Pf*A-M1. These results demonstrate that residues within the peptide substrate play an integral role in substrate recognition and binding, which ultimately influences the efficiency with which peptide substrates are processed.

The *Plasmodium* M17 aminopeptidases were conserved in their preference for large and non-polar residues in the P1 position such as Leu, Tyr, Trp and Phe. The limited capacity for residues in the P1 position is primarily due to the hydrophobic nature of the active site, and although the crystal structures of *Pb*-M17 and *Pv*-M17 are not yet available, the high degree of sequence conservation in these regions likely means the hydrophobic nature is also conserved in these two homologs. The *Plasmodium* M17 aminopeptidases have a highly limited specificity fingerprint when compared to homologs from porcine kidney (pkLAP), tomato (LAP-A), *E. coli* (PepA), *Helicobacter pylori* (*Hp*M17AP) and *Staphylococcus aureus* (*Sa*M17AP) which all show capacity to process a wider range of residues in the P1 position, such as Arg and Ala (17, 32, 33). The ability to process substrates with Arg in the P1 position is associated with a more hydrophilic active site cavity, as seen with *Hp*M17AP and *Sa*M17AP, and the inclusion of a sodium ion at the base of the helix connecting the C- and N-terminal domains (32, 33).

The M17 aminopeptidases demonstrated processive cleavage of the Leu-based substrates and, unlike the M1 aminopeptidases, catalytic behaviors were mostly conserved between homologs. As substrate length increased, we observed a conserved reduction in catalytic efficiency, which was largely due to a steady decease in substrate turnover rates. As all six active sites of the biologically active hexamer line the same internal cavity, a reduction in substrate turnover rates may be a function of longer peptide substrates occluding neighboring active sites. By occluding neighboring active sites, there would be fewer sites available to hydrolyze substrate, therefore slowing overall substrate turnover rates. The idea of increased interaction between active sites is supported by the increase in cooperativity as the substrate length increases. The three M17 homologs did not exhibit cooperativity when processing the single Leu residue but become increasingly cooperative as substrate length increased. Most M17-family aminopeptidases share this behavior and do not show evidence of allostery or cooperativity when hydrolyzing single residues linked to fluorescent reporters. An exception to this is the allosteric behavior of *Hp*M17AP, which has clear cooperative activity upon hydrolysis of Leu-p-NA (Dong, 2005). As little research has been conducted on M17 kinetics in the presence of longer peptide substrates, we cannot conclude if increased active site cooperativity due to increased substrate length is a conserved kinetic characteristic amongst the M17 aminopeptidase family or something unique to *Plasmodium* M17 aminopeptidases.

Activity levels against the Ala screen peptides compared to the hemoglobin-derived hexapeptide (LDPENF) varied between homologs, however the overall preference pattern remained conserved. *Pv*-M17 and *Pb*-M17 both had a low tolerance for Ala in the P4’ position in place of Asp, whereas *Pf*A-M17 tolerated Ala and Asp similarly in the P4’ position. The reason for this behavioral difference is unclear, as the location of the S4’ binding site within M17 aminopeptidases is unknown. Further to this, the known substrate sites (S1 and S1’) are highly conserved, as are the surrounding regions where the downstream binding pockets may be located. *Pv*-M17 and *Pb*-M17 preference for Asn in the P4’ position may be more related to substrate access to the active site through the N-terminal channels rather than substrate recognition and binding in the substrate binding sites. The three homologs shared a low tolerance for Ala in the P1, P2’ and P5’ position. The low activity against ADPENF was unsurprising, as the M17 aminopeptidases were shown to have low or no activity against Ala-ACC in the P1 screens. The larger size of the ADPENF peptide in relation to the small Ala-ACC substrate used in the P1 screen may help to anchor the peptide substrate in the active site and facilitate catalysis. The three homologs demonstrated no activity against LDPENA within the time period of this experiment. This peptide substrate is hydrolysable; we saw in earlier *Pv*-M17 and *Pb*-M17 time trials that the substrate can be cleaved, albeit at a very slow rate compared to the other screen peptides (Supp Fig S3). M17 aminopeptidases have highly hydrophobic and buried active sites, which may both require a level of hydrophobicity at the peptide C-terminal to successfully interact with and bind the substrate binding sites.

There were no other substitutions that resulted in notable differences in activity. This could be that M17 aminopeptidases are not particular in the residues accepted at certain subsites, or that Ala is also well-accepted in these positions. Further characterization of subsite specificities would provide invaluable insight into the functionality of M17, and M1, aminopeptidases and would be a welcome addition to the profiles outlined in this study.

## Conclusions

In this study, we have shown that the M1 and M17 aminopeptidases from three *Plasmodium* species display largely conserved substrate specificity profiles and kinetic behaviors within their families. We demostrated that *Plasmodium* M1 and M17 aminopeptidases change their kinetic behaviors when hydrolyzing substrates of different lengths and have distinct preferences for residues at various intra-peptide sequence positions, not just at the N-terminal. We can use this knowledge to broaden our understanding of the M1 and M17 aminopeptidase families and more specifically use our finding to anticipate challenges in designing inhibitory compounds that are selective, cross-species and cross-family targeting.

## Associated Content

### Supporting Information

**Supplementary Figure S1.** Representative mass spectrometry trace of LDPENF digestion by *Pb*-M17 (PDF)

**Supplementary Figure S2.** Digestion of Ala screen peptides by *Pv*-M1 and *Pb*-M1 in 45-min incubation period (PDF)

**Supplementary Figure S3.** Digestion of Ala screen peptides by *Pv*-M17 and *Pb*-M17 in 45-min incubation period (PDF)

### Accession Codes

*Pf*A-M1, UniProt O96935; *Pf*A-M17, UniProt Q8IL11; *Pv*-M1, UniProt A5JZA9; *Pv*-M17, UniProt A5K3U9; *Pb*-M1, UniProt A0A509ATR4; *Pb*-M17, UniProt A0A509APA2.

### Author Contributions

TRM performed experiments, analyzed data, co-wrote paper; BKH, MD, KWS and ND performed experiments, analyzed data; SM performed experiments, analyzed data, co-wrote paper and project concept

### Funding Sources

This work was supported by the National Health and Medical Research Council (Synergy Grant 1185354 to SM). The Drag laboratory is supported by National Science Centre in Poland and “TEAM/2017-4/32” project, which is carried out within the TEAM program of the Foundation for Polish Science, co-financed by the European Union under the European Regional Development Fund.

## Acknowledgements

TRM and BKH are supported by RTP scholarship from Australian Department of Education and TRM by the Monash Graduate Completion Award.

## Abbreviations

LAP: Leucine Aminopeptidase
ACC: 7-amino-4-carbamoylmethylcoumarin
NHMec: 7-amido-4-methyl-coumarin
Leu-Mec: Leucine-7-amido-4-methyl-coumarin

## Notes

### Competing Interest Statement

The authors have declared no competing interest.

