## Supplementary material for "Mapping the substrate sequence and length of the *Plasmodium* M1 and M17 aminopeptidases": Supp Fig S1

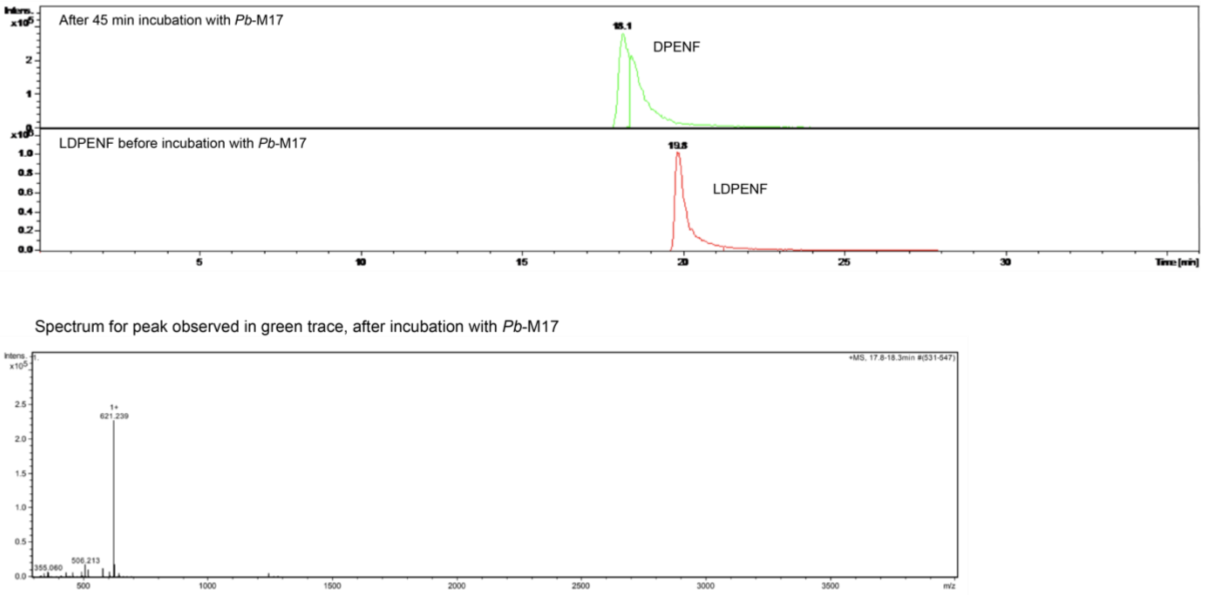


**Supplementary Figure S1. Representative mass spectrometry trace of LDPENF digestion by *Pb*-M17**. After 45 mins of incubation, only one product at a molecular weight of 621 Da is observed. This correlated to the removal of single peptide Leu from the N-terminal of hexapeptide LDPENF, producing pentapeptide DPENF. Trace is representative of cleavage results obtained for each aminopeptidase – in each experiment only peptides of molecular weights 734 Da (LDPENF) and 621 Da (DPENF) were observed.
