## Supplementary material for "Mapping the substrate sequence and length of the *Plasmodium* M1 and M17 aminopeptidases": Supp Fig S2

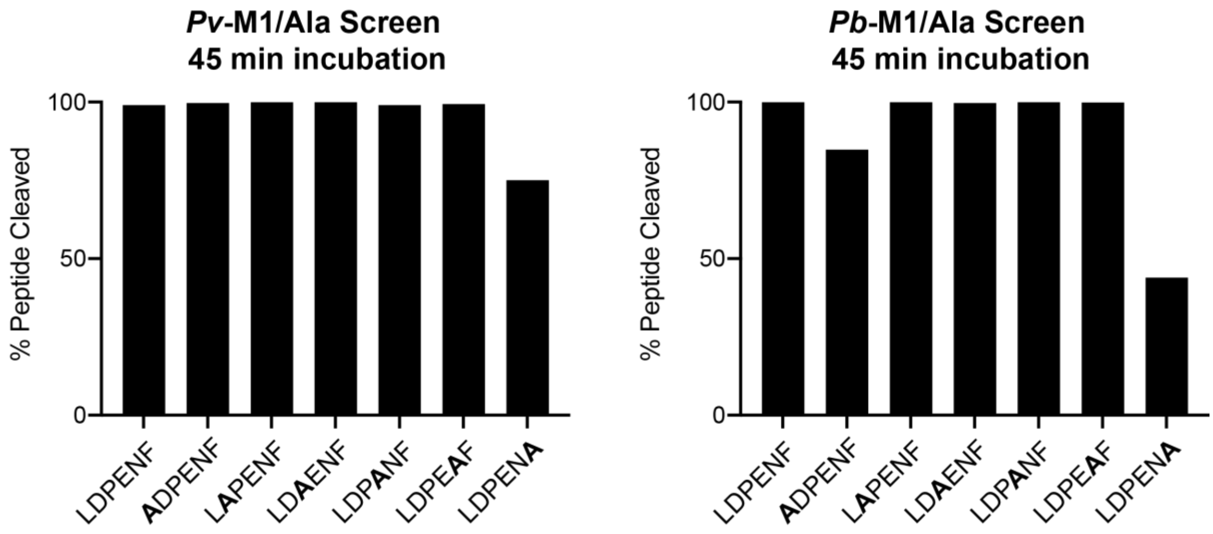


**Supplementary Figure S2. Digestion of Ala screen peptides by *Pv*-M1 and *Pb*-M1 in 45-min incubation period.** Template hexapeptide (LDPENF) and Ala screen peptides were digested to almost 100% in the 45-min incubation period at 37˚C. *Pv*-M1 and *Pb*-M1 showed decreased activity levels against peptides ADPENF and LDPENA
