## Supplementary material for "Mapping the substrate sequence and length of the *Plasmodium* M1 and M17 aminopeptidases": Supp Fig S3

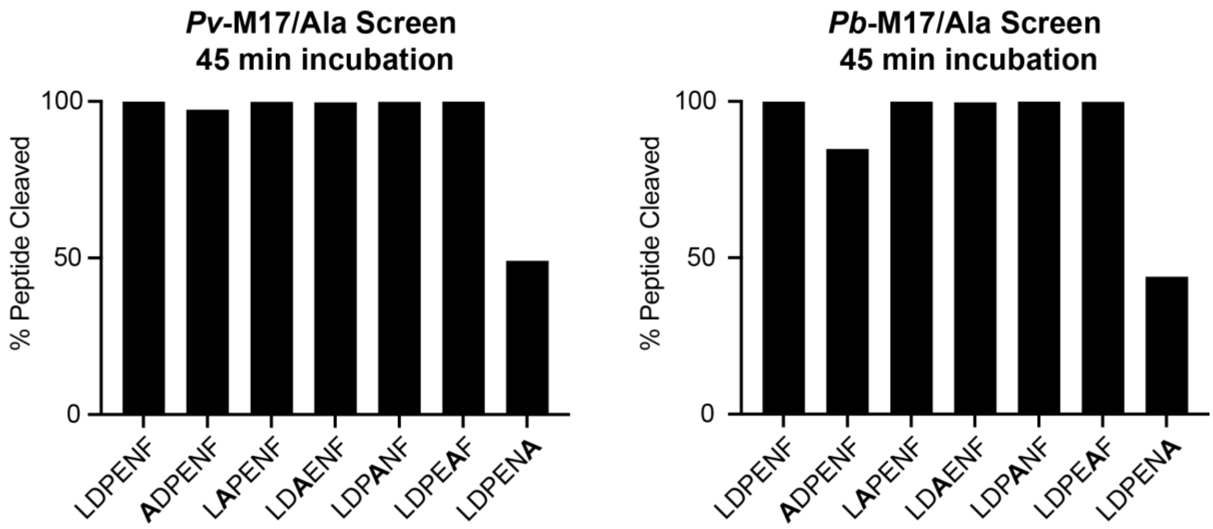


**Supplementary Figure S3. Digestion of Ala screen peptides by *Pv*-M17 and *Pb*-M17 in 45-min incubation period.** Template hexapeptide (LDPENF) and Ala screen peptides were digested to almost 100% in the 45-min incubation period at 37˚C. *Pv*-M17 and *Pb*-M17 showed decreased activity levels against peptides ADPENF and LDPENA
